# High-Resolution 3D Fluorescent Imaging of Intact Tissues

**DOI:** 10.1101/855254

**Authors:** Danny El-Nachef, Amy M. Martinson, Xiulan Yang, Charles E. Murry, W. Robb MacLellan

## Abstract

Histological analysis of fluorescently labeled tissues has been a critical tool to understand molecular organization *in situ*. However, assessing molecular structures within large cells and in the context of human organ anatomy has been challenging because it requires penetration of staining reagents and light deep into opaque tissues, while also conforming to the spatial constraints of high-resolution objective lenses. This methodology article describes optimized sample preparation for sub-micron resolution 3D imaging in human and rodent tissues, yielding imaging depth (>100 µm) and resolution (<0.012 µm^3^ voxel size) that has previously been limited to whole-mount *in vitro* organoid systems, embryos, and small model organisms. Confocal images of adult human and rodent organs, including heart, kidney, and liver, were generated for several chemical and antibody stains in cleared tissue sections >100 µm thick. This method can be readily adopted by any lab performing routine histology and takes 3 days from the start of tissue preparation to 3D images.

## Introduction

Analysis of tissue sections has been essential to our understanding of the molecular and cellular mechanisms that control structure and function in mammalian tissues and organs. In fact, the most common histological protocols such as hematoxylin staining of formalin fixed paraffin-embedded microtome sections have changed little since their development in the 1800’s^1^. The ability to generate and analyze thin tissue sections (i.e. <10 µm thick) *in situ* broadly transformed the field because it enabled minimal light scattering, efficient staining reagent penetration, and compatibility with high resolution objective lenses that have small working distances. While fluorescent molecular labeling and imaging modalities have continued to improve resolution of fine structures in 3D, the ability to study these structures within the context of cells or anatomical features that are >10 µm large is limited by light scattering, staining reagent penetration, and high-resolution objective lens working distances^2^. Several tissue clearing approaches have overcome light scattering, but the latter limitations remain^2^.

Tissue-clearing protocols that have attempted to address light scattering and penetration limits were recently reviewed^2, 3^. While most of these approaches were restricted to embryos and small animal models, three studies demonstrated tissue clearing in human brain or prostate samples^4–6^, though it is unknown how these protocols perform in other human organs. The major limitation shared by all previous reports is that the cleared samples were not imageable with high resolution objectives^6–8^. For the samples to physically fit within the working distance of the objective lens, low powered objectives (i.e. 20x dry, 0.6 numerical aperture (NA)) with large working distances (>500 µm) were used. These low NA objectives have limited ability to resolve fine structures because the resolution limit of an objective lens is determined by its NA^7, 9^. Abbe’s law defines the lateral (XY) resolution limit as λ/2NA; while axial (XZ and YZ) resolution is defined by 2λ/(NA^2^). Thus, objectives with lower NA have poorer lateral resolution and exponentially worse axial resolution. Recently designed objectives have been developed for cleared tissues that have large working distances and improved NA, but these require custom manufacturing and only modestly improve the NA to 0.9^10^. Artificial expansion of specimens can improve spatial resolution with standard large working distance, low NA objectives, but this requires crosslinking proteins to expandable hydrogels^8^, which can introduce distortions from uneven expansion^11^ and dramatically increases sample preparation and imaging time. All reported tissue clearing protocols, even those that do not require expansion microscopy techniques, have practical limitations because the procedure can take several weeks. For example, the Advanced CLARITY protocol (based on CLARITY) takes 7-28 days to perform^7^ and the iDISCO protocol (based on 3DISCO) takes 8-18 days^12^. Furthermore, many of these studies were restricted to examination of endogenously encoded fluorescent proteins and chemical stains, as antibody-based stains penetrate less efficiently^13–15^. While these approaches can decrease sample preparation time, they require the generation of transgenic lines for each molecule to be studied^13^. In addition, studies in cleared tissues have frequently utilized two-photon excitation fluorescence microscopy to fully utilize the working distance of low resolution objectives, extending imaging depth to several millimeters; but this requires costly, specialized equipment, and has limitations for multichannel acquisitions^16, 17^. For instance, DNA dyes and fluorescent proteins that can normally be distinguished by excitation wavelength in single photon microscopy demonstrate overlap with two-photon excitation^16–18^. Another practical consideration is clearing protocols often utilize customized electrophoresis chambers^7^ or proprietary reagents^2, 19^ that can be inaccessible or prohibitively expensive for analysis of large specimens. Recently, a protocol was established to clear and image organoids, enabling high resolution 3D imaging of human tissues^20^ but these engineered tissue constructs are much smaller in scale compared to adult mammalian organs and may not completely reflect matured cells within adult tissues *in vivo*. Advances in micro computed tomography have enabled imaging of thick human specimens, but this modality is not compatible with detection of fluorescent labels and cannot achieve the same resolution as optical approaches^21, 22^. While electron microscopy approaches have superior resolution, they are not suitable for 3D imaging in thick specimens^7^.

These challenges prompted us to develop a sample preparation and imaging technique that generates 100-200 µm thick sections in suspension, enabling rapid staining and clearing of samples that are compatible with standard high-resolution objectives (i.e. 60x oil, NA=1.4) and single photon confocal microscopes. These features yielded submicron resolution in all three dimensions within thick tissues that could be imaged in their entirety. Our use of defined, inexpensive clearing reagents and common lab equipment makes this approach accessible to any lab that performs routine histology. Together, this 3-day technique enables assessment of molecular structures in a variety of thick human biopsies and organs, such as cardiac myocyte sarcomeres, gap junctions, and T-tubules, hepatocyte canaliculi, kidney podocyte membranes, capillary networks, and subnuclear chromatin domains.

## Results

### Development of the novel histology plus clearing technique for submicron resolution in 3D

Empowering immunolabeling and high resolution imaging techniques in thick tissues has special relevance to cardiology and studies of cardiac myocytes (CMs) which are >100 µm in size^23^. The inability to resolve small macromolecules, such as Aurora Kinase B localized to cytokinesis cleavage furrows or condensed chromatin labeled with mitosis marker phosphorylated histone H3, in whole CMs *in situ* has led to contentious views on the capacity of adult mammalian CMs to proliferate^24–26^. Dissociation of the heart into single cells and subsequent *in vitro* analysis may bias against cells sensitive to protease digestion and might not reflect CM behaviors *in vivo*^27^, and identification of which cell types are expressing cell proliferation markers *in situ* has been equivocal using current histology methods that fail to capture CMs in their entirety or clearly resolve individual cells within the densely packed myocardium^24^. Thus, a sample preparation and imaging technique that provides volumetric quantifications and resolves sarcomere structure, chromatin organization, nuclei number, and cell cycle activity *in situ* in whole adult mammalian tissue holds great promise to address these intractable problems.

To this end, we used protocols we established for whole-mount high resolution 3D imaging of cleared embryos^28, 29^ as a starting point. Tissue fixation, embedding, sectioning, staining, optical clearing, and imaging protocols were tested to optimize a procedure in adult mammalian tissues for generating thick, optically clear specimens that could be imaged with sub-micron resolution in 3D. We found that lightly crosslinking the samples and post-fixation treatment with methanol was critical to maintain tissue integrity and for efficient downstream tissue clearing (**Figure 1a**). Agarose embedding, vibratome sectioning, and benzyl-alcohol benzyl-benzoate (BABB) solubilization effectively generated >100 µm thick, optically clear samples (**Figure 1a**) that could be imaged in their entirety with high-powered objective lenses and standard single-photon confocal microscopy.

**Figure 1.**
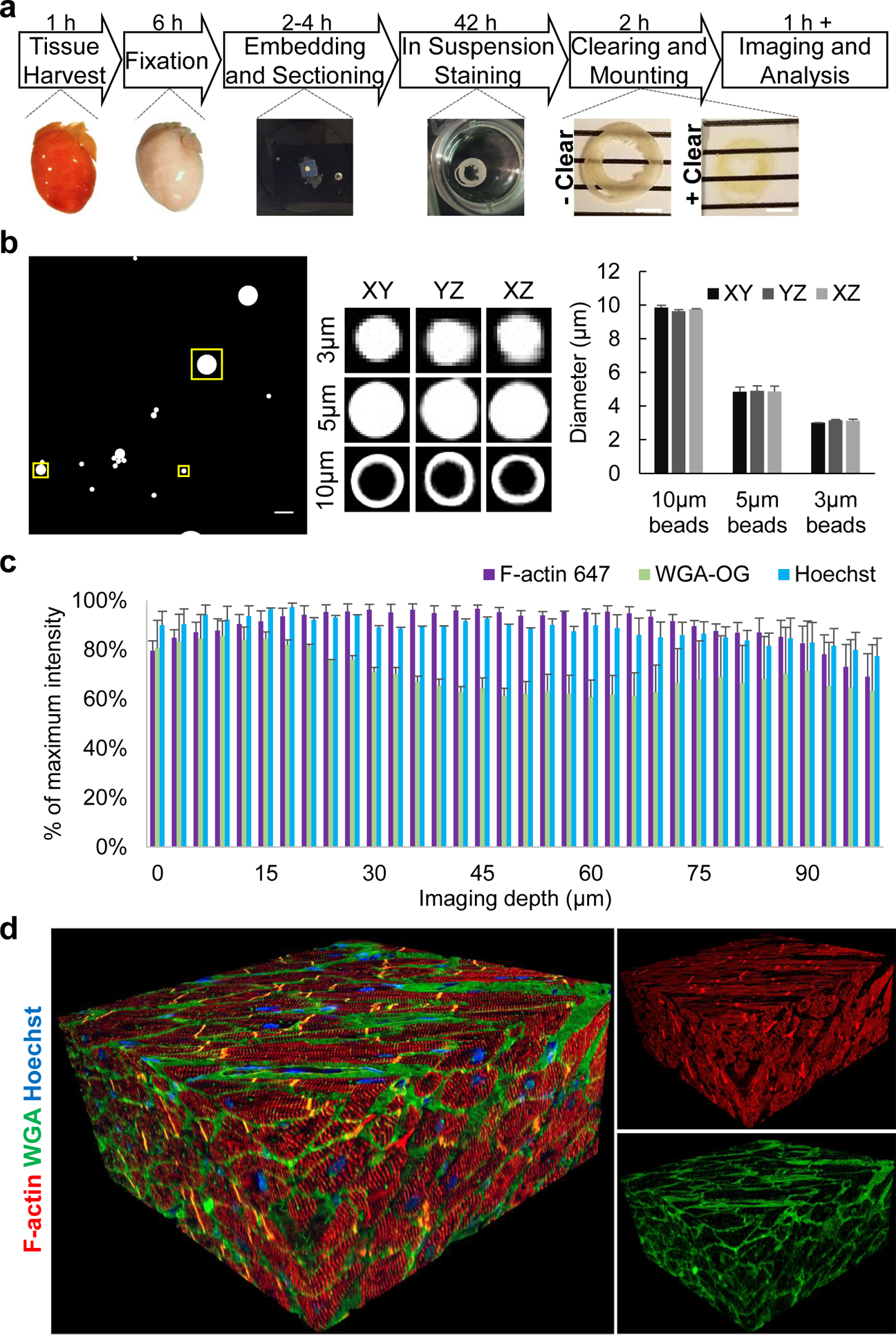
Overview of method and quality control. (**a**) Overview of protocol showing major steps and timing from tissue harvesting to imaging. Scale bar=4 mm. (**b**) Quality control representative images of fluorescent microbeads of different sizes mounted in BABB with a #0 coverslip, as done with subsequent tissue samples. The large image is a maximum intensity image of the z-stack with the boxed regions being 3.75, 6.26, and 12.73 µm^2^ and enclosing 3, 5, or 10 µm beads, respectively. Note the 10 µm beads are only fluorescently labeled at the bead surface. The boxed regions were examined with orthogonal slices of the z-stack and the maximum diameter slice for each dimension are shown in representative images and quantified. (**c**) Quality control for signal intensity through 100 µm of cardiac tissue using Z-intensity profiling. The average signal intensity per z-slice confirmed uniform staining and clearing for various fluorescent stains. (**d**) Representative 3D reconstruction of adult normal human heart. F-actin (red), WGA (green), and Hoechst (blue) staining from 373 confocal images, z-step size=300 nm. Acquired with 60x oil NA=1.4 objective.

Prior to 3D imaging the tissue samples, fluorescent beads manufactured with known diameters were examined using the same slide preparation and BABB mounting media as for the tissues to ensure lack of spherical aberrations, z-stretching, or z-compaction (**Figure 1b**). Normal adult human heart tissues were processed and stained for several markers, and Z-intensity profiles were plotted, demonstrating efficient clearing, even staining through the tissue depth, and acceptable levels of photobleaching (**Figure 1c**). Visualization of 3D reconstructions provided a qualitative assessment of the method’s performance (**Figure 1d**).

### Definitive assessment of cell cycling and chromatin in intact human and mice cardiac myocytes

To assess the proliferative capacity of human adult CMs we subjected normal and cardiomyopathy specimens to this technique and immunostained for a chromatin marker of mitosis, phosphorylated histone H3 (pH3). We found no evidence of adult CM proliferation in normal (**Figure 2a** and **Videos 1 and 2**) or cardiomyopathy (**Figure 2b**) human samples, nor in wildtype mice (**Figure 2c** and **Video 3**), though mitosis marker pH3 was unequivocally detected in cardiac myocyte nuclei from transgenic mice where the Myc oncogene is specifically induced in adult CMs^30^ (**Figure 2d** and **Video 4**). Importantly, the images demonstrated this method can be used to unequivocally assign nuclei as belonging to CMs or non-CM cell types, and could resolve fine molecular structures, such as individual sarcomeres, T-tubules, and capillaries, in all three dimensions within whole CMs in thick cardiac tissues. Because each z-stack is composed of hundreds of serial images that can take over an hour to acquire on a standard single-photon confocal, we tested this sample preparation with spinning disk confocal microscopy (**Figure 2e**). This demonstrated comparable 3D reconstructions could be generated in one tenth of the time as standard confocal microscopes and that the sample preparation method is compatible with multiple imaging modalities.

**Figure 2.**
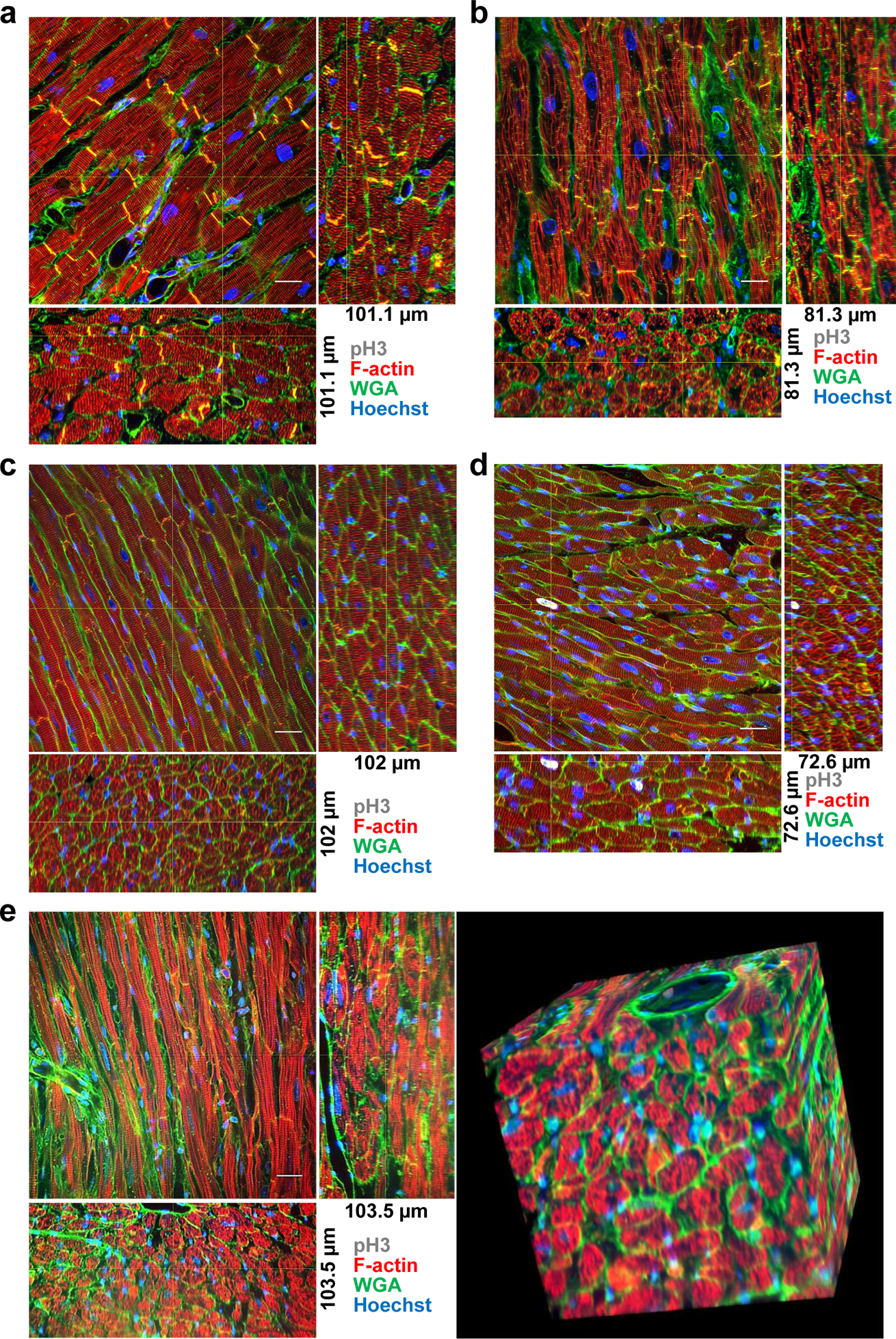
Adult cardiac myocyte cell cycling. (**a**) Normal human, (**b**) human cardiomyopathy sample from left-ventricular assist device core biopsy, (**c**) wildtype mouse, and (**d**) cardiac myocyte specific Myc-activation transgenic mouse samples stained with F-actin (red), WGA (green), Hoechst (blue), and pH3 (white). Orthogonal slices, indicated by the yellow cross-lines, from 3D reconstructions are shown. (**e**) Spinning disk microscopy image of wildtype mouse sample. (**a**) 373 confocal images z-step size=300 nm, (**b**) 271 confocal images z-step size=300 nm, (**c**) 102 confocal images z-step size=1 µm, (**d**) 242 confocal images z-step size=300 nm, (**e**) 345 confocal images z-step size=300 nm. All images were acquired with 60x oil NA=1.4 objective.

Since epigenetic modifications on chromatin and nuclear morphology have been identified as key mediators of CM proliferation and cardiomyopathies^23, 31–33^, we tested whether these features could be resolved by our technique. Nuclear morphology and subnuclear heterochromatin domains that stain positive for canonical heterochromatin markers Heterochromatin Protein 1 (HP1) and histone H3 Lysine 9 trimethylation (H3K9me3)^23^, were clearly resolved in 3D (**Figure 3**).

**Figure 3.**
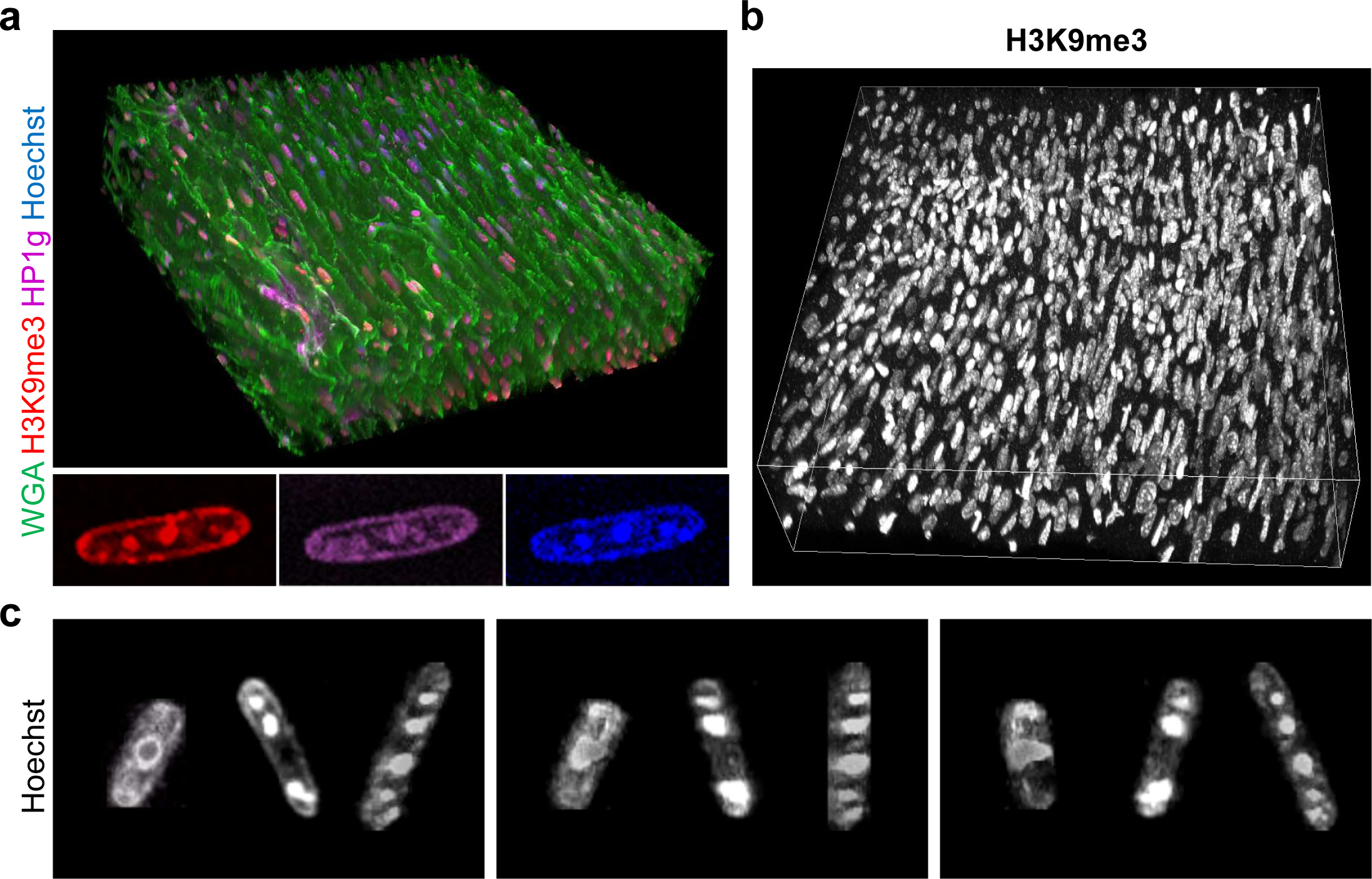
Adult cardiac myocyte heterochromatin and subnuclear chromatin domains. (**a**) Wildtype mouse cardiac sections were stained with WGA (green), Hoechst (blue), and heterochromatin markers histone H3 lysine 9 trimethylation (H3K9me3, red) and Heterochromatin Protein 1 gamma (HP1g, magenta). Insets, a single cardiac myocyte nucleus cropped from the z-stack shows colocalization of H3K9me3 and HP1g at subnuclear heterochromatin domains. (**b**) H3K9me3 (white) staining in wildtype mouse cardiac sample. (**c**) Three cardiac myocyte nuclei were cropped showing heterogeneous nuclear morphology. Each image represents a 90° rotation around the z-axis. (**a-b**) 270 confocal images z-step size=300 nm. All images were acquired with 60x oil NA=1.4 objective.

### Engrafted human pluripotent stem cell-derived cardiac cells partially integrate with host hearts

We next tested whether this method could enable assessment of cardiac cell therapy engraftment and integration with host myocardium at the molecular level in 3D. GFP-labeled human pluripotent stem cell-derived cardiac myocytes were injected into immune-compromised host rat myocardium as previously described^34, 35^ (**Figure 4**). The cleared thick sections were visualized with low-powered objectives to examine the total graft size in relation to total host myocardium, and with high-powered objectives that could resolve sarcomeres in the engrafted myocytes (**Figures 4a** and **4b**). As expected, no GFP+ cells were detected in vehicle injected rat hearts, and Connexin 43+ gap junctions in the rat myocardium were resolvable in 3D (**Figure 4c** and **Video 5**). Analysis of cell-injected animals revealed Connexin 43+ gap junctions connecting the engrafted cells and host myocardium 6-weeks after cell transplantation, which was confirmed in all dimensions (**Figures 4d** and **4e**). The 3D reconstructions also demonstrated large regions of graft encapsulated in WGA+ glycoproteins, consistent with fibrosis^36^ (**Figures 4d** and **4e**). We have previously found fibrosis is resolved three months after engraftment into non-human primate hearts, but this physical barrier may hinder electrical integration with the host and contribute to the rare, transient arrhythmogenic potential of cardiac cell therapy grafts seen soon after cell transplantation^35, 37^.

**Figure 4.**
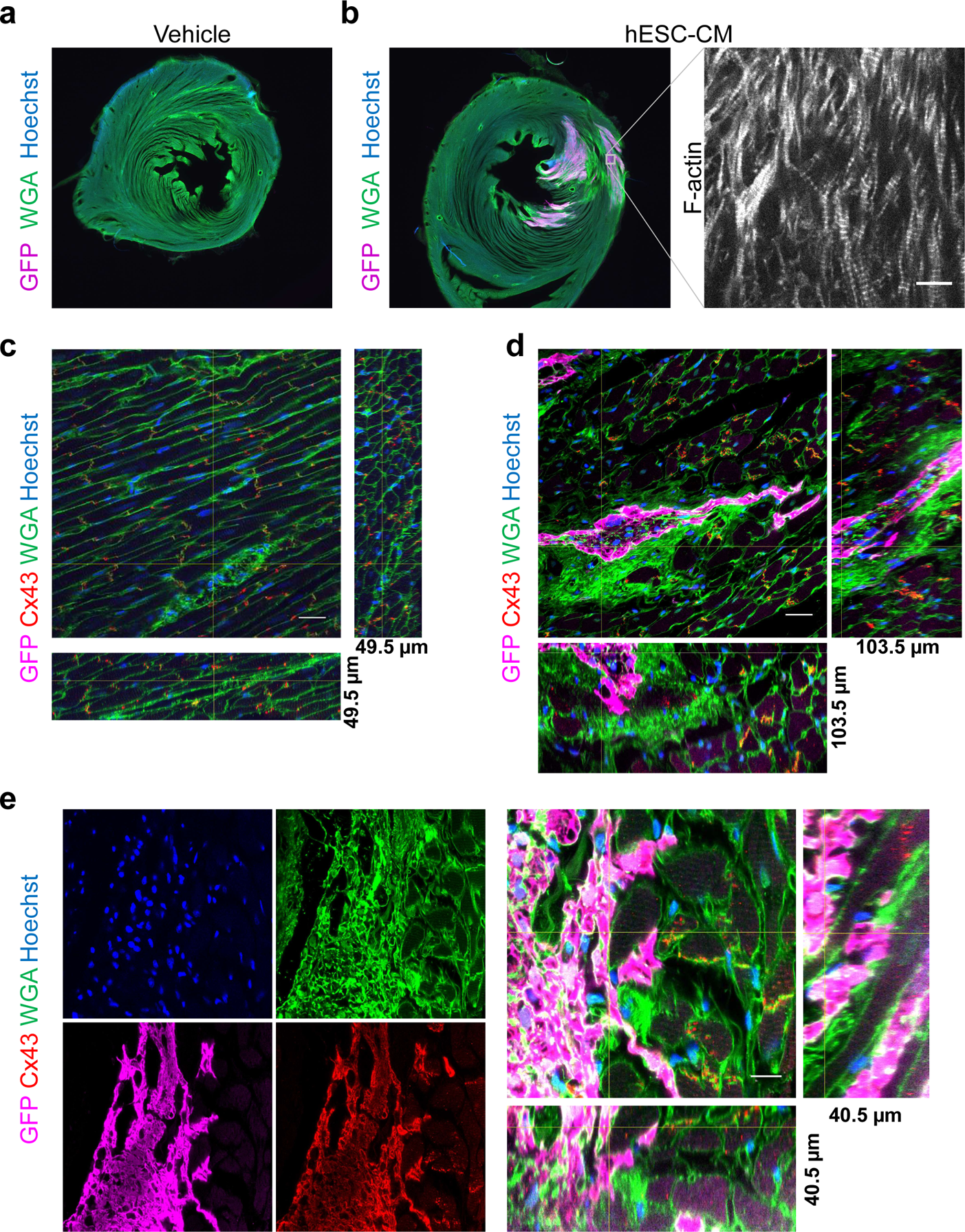
Cardiac cell therapy engraftment and myocyte gap junctions. Low magnification widefield images of immune compromised (athymic, cyclosporin treated) adult rat cardiac sections 6-weeks after injection with (**a**) vehicle control or (**b**) GFP expressing human embryonic stem cells differentiated to cardiac myocytes (hESC-CM), stained with GFP (magenta) WGA (green) and Hoechst (blue). Inset shows F-actin staining of the same region of graft in an adjacent section, bar=10 µm. (**c**) Vehicle injected rat cardiac section stained for GFP (magenta), Connexin 43 (Cx43, red), WGA (green) and Hoechst (blue) showing gap junctions. 165 confocal images z-step size=300 nm. Bar=20 µm. (**d**) hESC-CM injected rat sections stained for GFP (magenta), Connexin 43 (Cx43, red), WGA (green) and Hoechst (blue) GFP+ graft and gap junctions demonstrating large patches of fibrosis around graft. 345 confocal images z-step size=300 nm. Bar=20 µm. (**e**) Example of well-integrated graft showing each stain individually and a 3D reconstruction of the graft-host interface. 135 confocal images z-step size=300 nm. Bar=10 µm. (**c-e**) Orthogonal slices, indicated by the yellow cross-lines, from 3D reconstructions are shown from images acquired with a 60x oil NA=1.4 objective.

### Large field imaging and morphological assessment in human and mouse kidneys

One apparent limitation of high-powered objectives with small fields of view and short working distances is that they could be unable to completely capture larger structures, such as human glomeruli. Using the same sample preparation technique, high resolution images were obtained in human and mouse normal adult kidneys showing clearly resolved vasculature, tubules, podocytes and nuclei. To image entire human glomeruli, which exceed 100 µm in size, a 40x oil objective (NA=1.3) was used and displayed modest reduction in resolution (**Figure 5a** and **Video 6**). Tiling (i.e. stitching) multiple fields of view also demonstrated larger regions can be imaged without compromising resolution (**Figure 5b** and **Videos 7-9**).

**Figure 5.**
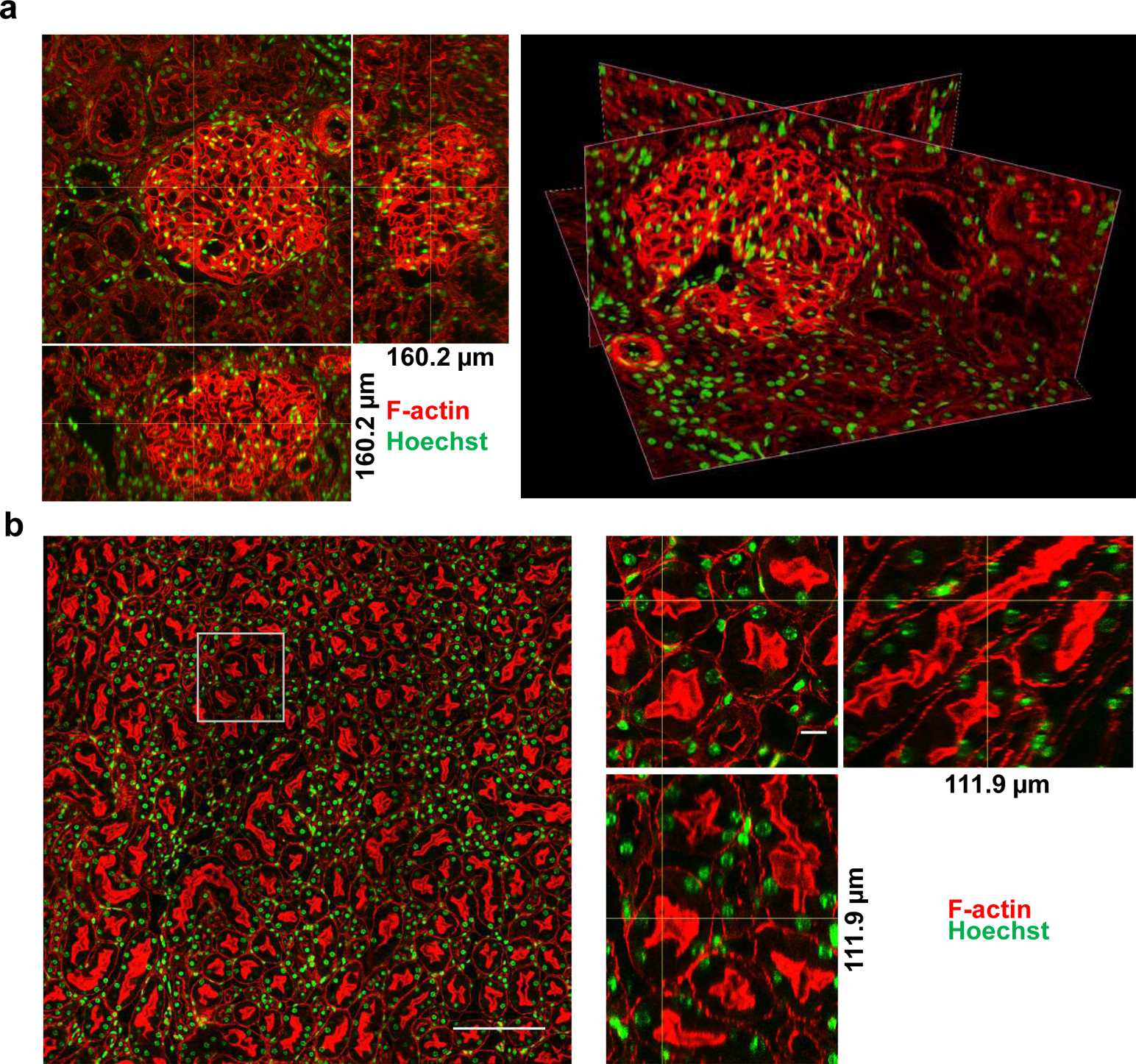
Adult kidney glomeruli and tubules. (**a**) Normal human orthogonal, left, and oblique slices, right, and (**b**) wildtype mouse orthogonal slices of kidney sections stained with F-actin (red) and Hoechst (green). (**a**) 534 confocal images z-step size=300 nm acquired with a 40x oil NA=1.3 objective. (**b**) 253 confocal images, z-step size=300 nm acquired with a 60x oil NA=1.4 objective with 4×4 fields of view tiled to examine larger area. Left, tiled image, bar=200 µm. Right, zoom in on boxed region in tiled image with orthogonal slices, bar=10 µm.

### Volumetric quantification of cells and subcellular structures in intact tissues

To demonstrate broader utility, we subjected liver sections from adult mice to analysis by this method (**Figure 6** and **Videos 9-11**). The resulting images were used for volumetric quantifications, such as hepatocyte volume, nuclei volume, and quantifying the number of nuclei per cell (**Figure 6**). For instance, we found that 72.9% of adult mouse hepatocytes were binucleated. This percentage closely mirrors previous *in vitro* studies in isolated hepatocytes^38^ but is substantially higher than other reports from *in situ* analysis that reported 27.5% binucleation^39^, which likely underrepresented the amount of binucleation due to the use of 3 µm thick sections that fail to capture whole hepatocytes. Binucleated hepatocyte volume was 64% larger compared to mononucleated hepatocytes (3614 ± 82.45 µm^3^ versus 2202 ± 84.63 µm^3^, P<0.0001, **Figure 6**), with the volume of individual nuclei showing no significant difference between these groups (159.6 ± 6.13 µm^3^ in mononucleated hepatocytes versus 150.3 ± 2.33 µm^3^ in binucleated hepatocytes, P<0.14). Consistently, the proportion of nuclear volume to cytoplasm volume was similar between the groups, with the fraction of total nuclear volume being 19% higher in binucleated hepatocytes. (0.081 ± 0.0028 in mononucleated hepatocytes versus 0.097 ± 0.0022 in binucleated hepatocytes, P<0.004, **Figure 6**).

**Figure 6.**
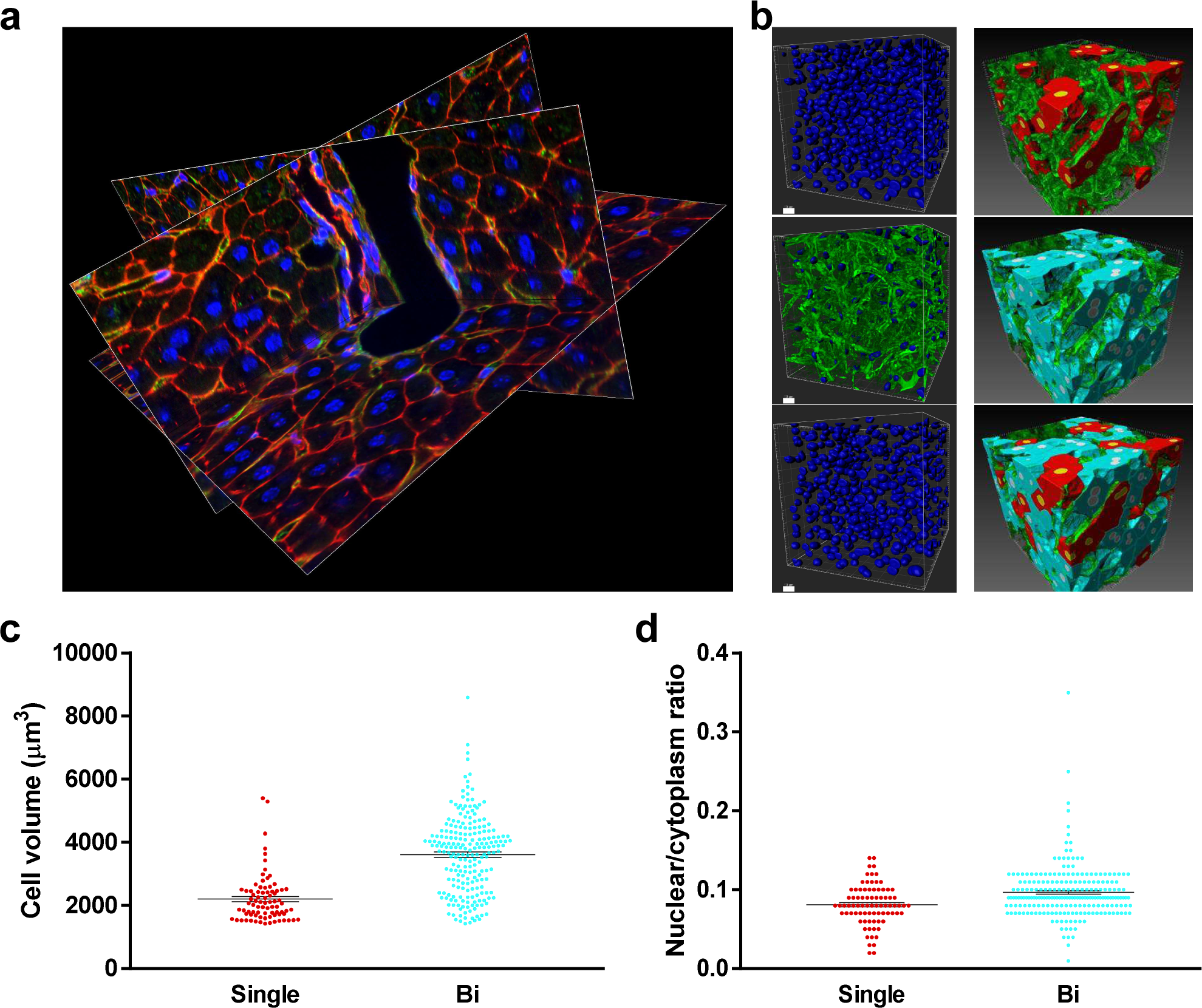
Adult liver vasculature and quantification of hepatocyte nuclei number. (**a**) Oblique slices of 3D reconstruction showing portal vein, bile duct, hepatocytes, vasculature, and nuclei stained with F-actin (red), WGA (green), and Hoechst (blue). 395 confocal images z-step size=300 nm. Acquired with 60x oil NA=1.4 objective. (**b**) Top left, segmentation of all nuclei. Middle left, nuclei colocalized with the vasculature. Bottom left, hepatocyte nuclei (nuclei from middle left were subtracted from top left), bars=10 µm. Right, segmentation of mononucleated hepatocytes (red), binucleated hepatocytes (turquoise), and vasculature (green). (**c**) Quantification of on mononucleated (single) and binucleated (bi) hepatocyte volume. (**d**) Total nuclear to cytoplasmic ratio in mononucleated and binucleated hepatocytes.

## Discussion

Here, we developed a combined histology and tissue clearing approach that is broadly applicable to studies of molecular and cellular three-dimensional organization in adult human and non-human mammalian specimens. Using common lab equipment and reagents, we show this method can be applied to studies of cardiac myocyte proliferation in adult human hearts samples; subnuclear chromatin architecture; cardiac myocyte gap junctions; human pluripotent stem cell-derived cardiac cell therapy engraftment in host rats; human kidney tubules, glomeruli, and vasculature; liver vessel networks and hepatocyte canalicular networks; and volumetric quantification of cells, subcellular structures, and nuclei number in large cells *in situ*. Using this technique, we determined that adult CMs are not proliferative in healthy or diseased human hearts. High resolution 3D imaging of human pluripotent stem cell (PSC)-derived cardiac myocytes engrafted in host rats demonstrated grafts integrate with the host myocardium but also revealed large portions of graft are encapsulated by fibrosis 6 weeks after implantation. Imaging of liver samples with intact hepatocytes demonstrated unprecedented levels of binucleation that correlated with cell hypertrophy. Interestingly, these images also revealed spiral-aligned nuclei in the liver bile duct epithelium and kidney tubules. We confirmed this method can be used with multiple imaging modalities including conventional single-photon line-scanning confocal and spinning disk microscopy. This methodology could be adapted for 3D pathological assessments of disease state and for generating submicron-resolution whole organ atlases. Indeed, subcellular features are emerging as key markers for disease progression, such as multinucleation, sarcomere and T-tubule density and alignment, capillary density and alignment, and canaliculi dilation^27, 40–47^. In addition, recent tissue engineering approaches demonstrated cells and extracellular matrix can be bioprinted with micron-level resolution^48, 49^, the datasets generated by this method will also be useful blueprints for engineering human tissues.

This method overcomes the limitations that have prevented high resolution 3D imaging of fluorescently labeled human biopsies, though, intrinsic limitations remain. For example, single photon confocal imaging illuminates the entire depth of the sample while imaging each plane, thus the sequential z-stack imaging could lead to photobleaching. Despite this we show this is not a limiting factor for most of the common fluorophores which have been designed with high extinction coefficients (i.e. Alexa Fluor conjugates and common DNA dyes)^50^. This issue could further be mitigated by two-photon imaging^51^. While this protocol maximizes the amount of specimen imageable with high powered objectives (i.e. 60x oil, NA=1.4), which are required for submicron resolution in 3D, going beyond 100 µm in depth is still hampered by the working distances of these objectives (210 µm), which is partially occupied by the coverslip, with even the thinnest coverslips (i.e. #0) being approximately 100 µm thick. This can be overcome partially by using lower NA objectives (i.e. 40x oil, NA=1.3) that have greater working distance, with modest reduction in resolution, enabling imaging of >160 µm of sample depth. This protocol quenches native fluorescence of genetically encoded fluorescent proteins. We overcame this by utilizing antibodies specific to green fluorescent protein (GFP), and such antibodies have been generated for all commonly used fluorescent proteins^52^. Lastly, the hundreds of z-stack images per specimen required for full utilization of the protocol can take over an hour to acquire and can be cumbersome to handle in image analysis software like ImageJ. We demonstrate these limitations can be ameliorated with faster imaging modalities (i.e. spinning disk confocal) and software (ie BitPlane Imaris, Nikon Elements) that enable faster acquisitions and analysis. Resonant scanning or light-sheet imaging are alternatives that could also increase acquisition speed^53^. Altogether this simple and rapid method achieves the highest levels of resolution and imaging depth that is possible using widely available standard confocal microscopy and broadly enables molecular biology and volumetric quantification studies in the context of three-dimensional human and mammalian tissues.

## Materials and Methods

The entire method is summarized in **Figure 1a** and the reagents and equipment used are listed in **Table 1**. Troubleshooting suggestions are provided in **Table 2**.

**Table 1.** Equipment and Reagents. List of equipment and reagents used in this study. * indicates hazardous reagent or equipment, please consult user manuals and MSDS for appropriate usage.

| Equipment |
| --- |
| 50mL tube rotator |
| Orbital shaker |
| Vibratome (Leica VT1200s) |
| *Vibratome blade (TED Pella, 121-9) |
| *Scalpel or surgical scissors |
| Sharps waste container |
| Confocal microscope (Nikon A1R) |
| PC with image analysis software (ImageJ/FIJI, Nikon elements, or BitPlane Imaris) |
| *Hot plate |
| Painbrush, soft, synthetic bristles, #0. |
| Blunt Forceps |
| Luer Lok syringe (optional, for perfusion) |
| Blunt canula (optional, for perfusion) |
| Canula clip (optional, for perfusion) |
| Suture (optional, for perfusion) |
| Cryomold |
| Lab tissue paper (i.e. Kimwipes) or gauss |
| Cotton swab |
| Nitrile gloves |
| Polypropylene tubes (1.5mL, 5mL) |
| Nylon Mesh or Cell Strainer |
| 24 well plates |
| Falcon tubes (50mL) |
| Sample baskets (5mL tube + cell strainers/mesh) |
| Petri dishes |
| Coverslips (#0) |
| 100mL Glass Jars |
| Coverslip caddy |
| Slides |
| Adhesive, solvent-resistant foil (Nashua Foil Tape, 279 µm thickness, catalog# 1207802) |
| Reagents |
| Phosphate buffered saline (PBS) (Gibco catalog # 21300-58) |
| KB buffer (mmol/L: KCl 20, KH<sub>2</sub>PO<sub>4</sub> 10, K<sup>+</sup>-glutamate 70, MgCl<sub>2</sub> 1, glucose 25, taurine 20, EGTA 0.5, HEPES 10, 0.1% bovine serum albumin, pH adjusted to 7.4 with KOH) |
| *16% paraformaldehyde, ampoule in nitrogen. (Alfa Aesar catalog # 43368) |
| *Methanol (Sigma catalog# 34860) |
| Low melt agarose (Fisher Scientific catalog # BP165-25) |
| Bovine Serum Albumin (BSA) |
| *Dimethyl Sulfoxide (DMSO) |
| *Sodium Azide |
| *Super glue |
| Normal Donkey Serum |
| *Triton X-100 |
| Fluorescent beads (Fluorsbrite, Thermofisher, Sigma) |
| Primary antibodies: phosphorylated serine 10 histone H3 (pH3, Abcam ab5176), trimethyl lysine 9 histone H3 (H3K9me3, Active Motif 39161), Heterochromatin Protein 1 gamma (HP1g, Active Motif 39981), Connexin 43 (Cx43) (Zymed catalog # 71-0700), GFP (Novus catalog # NB100-1770). |
| Secondary antibodies: (Alexa Fluor 488, 555, and 647, Thermo Fisher) |
| Chemical dyes: Wheat Germ Agglutinin (WGA) Alexa Fluor-647, WGA-Oregon Green, phalloidin-Alexa Fluor 647, phalloidin-rhodamine, Hoechst 33342 |
| Poly-L-lysine |
| *Isopropanol |
| *Benzyl alcohol |
| *Benzyl benzoate |
| Nail polish (Sally Hanson Insta-dry, clear) |

**Table 2.** Suggested Troubleshooting. Potential problems and solutions when using this method.

| Problem | Solution |
| --- | --- |
| Sample comes out of agarose block during section | Work quickly with the molten agarose, blot the tissue gently but thoroughly, swirl the tissue in the molten agarose so that any residual PBS is mixed in with the agarose. Ensure there is enough agarose surrounding the tissue after trimming the block and that the block remains cold during sectioning. |
| Morphology artifacts | Use soft brushes, careful handling, slower vibratome blade advancing. Adjust fixation times. Adjust agarose % for embedding. |
| Thickness of section not accurate | We confirmed the vibratome section thickness is calibrated using Leica Vibrocheck. If the user does not have access to a vibratome calibrator, they can check the thickness by performing z-stacks on a confocal microscope and adjust the vibratome settings to compensate for differences in visualized thickness versus expected thickness. When using confocal imaging to assess section thickness, it is essential that the user imaging configuration has been validated with fluorescent bead standards |
| Low signal (broadly) | Use cold PFA and fix for no longer than 3-6 hours. Ensure MeOH is not warming up during post-fix. Keep isopropanol incubations at 50 seconds each, extending this step will reduce signal. Use fresh samples or store samples in accordance with the protocol notes to prevent signal decay. |
| Low signal (interior/middle of section) | Add 1% DMSO to staining buffer, increase rocker speed, perform Ab staining steps at room temp, or extend incubation time. Increase staining reagent concentration. Though not suitable for studies quantifying signal intensity, uneven staining across the thickness can be overcome using adaptive imaging where excitation laser power is adjusted throughout the acquisition to give uniform signal, or post processing algorithms, such as ImageJ/FIJI (Menu: Images→Adjust→Bleach Correction→Histogram Matching). |
| Distortions in z-dimension (broadly) | Adjust correction collar on objective lens or adjust coverslip thickness and confirm with fluorescent beads; or (not recommended for quantitative studies) correct z-step size post processing using calibration values determine by fluorescent microsphere standards. |
| Distortions in z-dimension (one-sided) | This is consistent with going over the working distance limits of the objective, where the objective begins pushing up against the coverslip and z-step size is no longer accurate. This can also potentially damage the objective. Use thinner coverslips (i.e. #0) or switch from a 60x oil objective to 40x oil for more working distance. |
| Sample cannot get in focus on high power objective | Use thin #0 coverslips. Ensure that the sample is flat and well-adhered on the coverslip throughout the clearing steps, taking care to transfer the samples gently from each solution. Gently handle the slides after sealing with nail polish. Though the sample is sealed between the coverslip and slide, if it gets detached from the coverslip this will result in less sample imageable within the working distance of the objective. |
| Computer crashes | Reduce bit-depth, the human eye cannot distinguish more than 250 shades of gray, thus even 8-bit images are suitable for most applications. Alternatively, sub-sample regions of interest. Commercial software, such as Nikon Elements or Bitplane Imaris, can handle large file sizes more efficiently. |

### Human sample preparation

All human samples were approved by the Institutional Review boards of the University of Washington in Seattle. Fresh samples were collected in cold (4°C) PBS, or for heart tissue samples were collected in a high potassium buffer, KB buffer (110mM [K^+^]), to keep the myocytes arrested. Tissues were trimmed tissue into ∼6 mm^3^ cubes or smaller. Samples were then transferred to a 50mL conical tube with 30mL of fresh 4% paraformaldehyde (PFA) in PBS, precooled to 4°C and incubated 3-6 hours at 4°C in a rotator. The samples were then transferred into a new 50mL conical tube containing 30mL 100% methanol (MeOH), precooled to −20°C and incubated at −20°C for 1 hour. In a stepwise manner, samples were incubated in 30mL of precooled 80% MeOH/20% PBS, 60%MeOH/40%PBS, with each incubation for 30 minutes at −20°C. The samples were then washed 3x with precooled PBS and kept in PBS at 4°C until embedding.

### Rodent sample preparation

All animal protocols in this study were approved by the University of Washington Office of Animal Welfare (IACUC protocol # 4290-01) and conform to the Guide for the Care and Use of Laboratory Animals published by the US National Institutes of Health. All mice studies were performed in animals 8-12 weeks of age. Cardiac cell therapy injections into immunocompromised adult rats were performed as described^54^. We administered 10 units of heparin per gram of body weight by intraperitoneal injection, allowed it to circulate for 10 minutes to prevent blood coagulation, and then euthanized the animals via isoflurane overdose. Whole organs were dissected from animals and collect into iced Petri dishes containing 4°C PBS or KB buffer (hearts). Extraneous tissue was trimmed. We cannulated aorta (hearts), portal vein (livers), or renal artery (kidneys) with a blunt 26g needle, and secured the canula with a clip and suture for organ perfusion: We perfused 10mL of PBS or KB buffer (hearts), precooled to 4°C, with a syringe at a flow rate of ∼ 2mL/min to wash out blood cells. The syringe was then replaced with a new one containing 10mL 4% PFA in PBS, precooled at 4°C, and tissues were perfusion fixed at a flow rate of ∼ 2mL/min. From here we followed the preparation as described for human samples, starting from incubation with 4% PFA for 3-6 hours at 4°C in a rotator through methanol fixation and PBS washes.

### Tissue Embedding and Sectioning

6% low-melt agarose in PBS was prepared in a 50mL tube and dissolved by boiling the tube in a beaker containing water on a hotplate with the tube lid loose. The solution was cooled to 45°C in a water bath incubator. Fixed specimens were dabbed on a Kimwipe or gauze and placed in a cryomold that was then quickly filled with the molten agarose. Samples were left at room temperature to allow gelation and stored at 4°C until sectioning.

The agarose-tissue block was removed from the mold and the agarose was trimmed such that there was ∼4mm of agarose surrounding each side of the tissue, keeping the sample on ice. The trimmed agarose block was superglued to a vibratome sectioning platform. The vibratome sample chamber was then filled with PBS, precooled to 4°C. For sectioning, we found slow blade advancement and moderate amplitude to be crucial for good sectioning. For a Leica 1200S vibratome we used 0.14mm/min blade advance speed, 0.5mm amplitude, and a 15° cut angle. Section thickness was set to 100-200µm. We ensured the vibratome section thickness was calibrated using Leica Vibro-Check. Sections were collected with a soft, synthetic-bristle paintbrush and transferred to a 24-well plate containing cold PBS. Sections were stored up to two weeks at 4°C in PBS + 0.1% Na-azide + 1% bovine serum albumin (BSA), or for longer periods (we tested up to one year) at −20°C in 10% DMSO 90% PBS with 2% BSA.

### In-suspension staining

We utilized a convenient staining protocol we developed specifically for vibratome sections. In contrast to other sections, vibratome sections remain free-floating in suspension throughout the staining protocol to improve reagent penetration. We used 24-well plates filled with PBS or staining reagents and home-made baskets to hold the sections and minimize handling. The sample baskets are plastic cylinders that have a nylon mesh at one end, allowing the exchange of fluids from the sample within the basket and the buffers in the 24-well plate wells. To make these, we trimmed the cap and bottom from a 5mL polypropylene tube to make an open cylinder. We used nylon mesh (ie Cell Strainer with 100µm mesh) that was then placed a piece of aluminum foil on a hot plate and briefly pressed the plastic cylinder against the nylon mesh on top of the foil/hot plate to melt the mesh onto the tube edge. Other vessels may also be suitable, but it is important to stain the samples in suspension where reagents can flow through and make contact with all sides of the sample (in contrast to after being adhered to glass allowing only partial contact with reagents), as this improves the penetration of antibodies and dyes into the interior of the section. We prepared 24-well plates with the solutions below in sequential order, using 600uL of solution per well, and simply transferred the sample basket sequentially from well to well. Sections were transferred into sample baskets using a soft synthetic bristle paintbrush. Samples were washed with PBS + 0.1% TritonX100 (PBST) three times at room temperature. Blocking was performed with 5% normal donkey serum in PBST for 1 hour at room temperature on an orbital rocker. Plates were then transferred to 4°C and rocked on orbital shaker for an additional 30 minutes. Samples were incubated in primary antibodies diluted in blocking buffer supplemented with 1% DMSO at 4°C for 16 hours with rocking on orbital rocker. Samples were washed PBST four times, 5 minutes per wash, on orbital rocker at room temperature. In the last wash, plates were moved to 4°C on an orbital rocker for an additional 30 minutes. Samples were incubated with secondary antibodies + chemical stains (such as DAPI, Hoechst, phalloidin, or wheat germ agglutinin, WGA) at 4°C for 16 hours with rocking. Samples were then washed with PBST four times, 5 minutes per wash, on an orbital rocker at room temperature. Lastly, samples were kept in PBS until mounting.

### Clearing and Mounting

We prepared glass slides by adhering two pieces of ∼300µm-thick solvent-resistant aluminum foil tape such that they were spaced far enough apart to accommodate the sample fitting in between them, but close enough so that both sides of the coverslip could rest on the foil. These acted as bridges for the coverslips to lean on, preventing the samples from being crushed or squeezed. 22×22 mm coverslips were coated with 0.01% poly-L-lysine in PBS. Sections were transferred from the basket into a Petri dish containing PBS and then onto the center of the coverslip with a paint brush. Residual PBS was blotted off. The coverslips with samples were placed in a coverslip rack/caddy, and submerged into glass jars containing 100mL of following solutions sequentially, 50 seconds each incubation: 70% isopropanol, 85% isopropanol, 95% isopropanol, 100% isopropanol x 2, Benzyl alcohol and benzyl benzoate at a 1:2 ratio (BABB) x3. The samples were kept in the final BABB incubation for at least 10 minutes and then transferred to the glass slide, placing the coverslip between the two foil bridges. Excess BABB was blotted off and the slides were sealed with 3 coats of nail polish

### Imaging

All images were obtained using standard methods on a Nikon A1R confocal microscope, except for Figure 2e where a Nikon TiE equipped with Yokogawa spinning disk head was used and Figure 4a-b where a widefield Nikon TiE microscope was used. A 60x oil NA=1.4 objective was used for all images, except for Figure 4a-b where a 10x dry objective was used and Figure 5a where a 40x oil NA=1.3 objective was used. Unless noted otherwise in the figure legend, all z-stacks were acquired with a 60x oil objective and 300 nm z-step size, resulting in a voxel size of < 0.012 µm^3^. While our microscopy configurations did not require adjustments when using #0 coverslips, coverslip thickness or the correction collar on objective lenses could be adjusted to circumvent imaging aberrations if seen with the fluorescent bead standards^55^.

### Visualization and analysis

ImageJ/FIJI was used to generate orthogonal slices of the 3D reconstructed z-stacks (Menu: Images→Stacks→Orthogonal views). Z-intensity profiling was performed for each channel using Image J (Menu: Images→Stacks→Plot Z-axis profile). We used ImageJ, Nikon Elements, and BitPlane Imaris to generate 3D renderings. Imaris was used for generating videos of orthogonal slices through the tissue volumes. For analysis of hepatocyte volume and nuclei we used Imaris V9.2 and the Imaris *Surfaces* and *Cell* modules. Nuclei (Hoechst), hepatocytes (F-actin), and liver vasculature (WGA) were segmented independently. The segmented vasculature mask was applied to the nuclei channel and subtracted to remove non-hepatocyte nuclei. The remaining nuclei were masked and the number of nuclear masks per segmented hepatocyte was quantified.

## Supporting information

Video 1_Heart_normal adult human

Video 2_Heart_normal adult human_WGA and nuclei

Video 3_Heart_wildtype mouse_pH3

Video 4_Heart_Myc transgenic mouse_pH3

Video 5_Heart_adult rat_Cx43

Video 6_Kidney_Human_glomerulus_40x

Video 7_Kidney_Human_tiled

Video 8_Kidney_mouse_tiled

Video 9_Liver_mouse_tiled

Video 10_Liver_mouse

Video 11_Liver vessels_mouse

## Acknowledgements

We would like to acknowledge April Stempien-Otero and Chris P. Miller for providing human samples. We are grateful to Dale Hailey and the University of Washington’s Garvey Imaging Core for microscopy support. We thank William C. Smith and Michael T. Veeman for the development of whole-mount embryo clearing protocols and consultation. This work was supported by AHA 14PRE20140025 and NIH F32HL143851 (DE), NIH R01 HL70748 (WRM), and by NIH grants R01HL128362, U54DK107979, R01HL128368, R01HL141570, R01HL146868, and grants from the Foundation Leducq Transatlantic Network of Excellence and Robert B. McMillen Foundation (CEM).

## Contributions

D.E. designed the study, performed experiments, analyzed results, and wrote the manuscript. A.M.M. performed cardiac cell therapy surgeries and animal care. X.Y. generated PSC-derived cardiac myocytes for cell therapy. W.R.M., C.E.M., and X.Y. edited the manuscript. W.R.M. and C.E.M. provided supervision and acquired funding.

---

**Video 1 Normal adult human heart.** F-actin (red), WGA (green), and Hoechst (blue) staining 3D reconstruction from 373 confocal images, z-step size=300 nm. Acquired with 60x oil NA=1.4 objective.

**Video 2 Normal adult human heart**. 3D reconstruction of the same images in Supplementary Video 1, with F-actin signal omitted to highlight T-tubules. Acquired with 60x oil NA=1.4 objective.

**Video 3 Wildtype adult mouse heart**. F-actin (red), WGA (green), Hoechst (blue), and pH3 (white) staining 3D reconstruction from 102 confocal images, z-step size=1 µm. Acquired with 60x oil NA=1.4 objective.

**Video 4 Transgenic adult mouse heart with adult cardiac myocyte specific activation of the Myc oncogene.** F-actin (red), WGA (green), Hoechst (blue), and pH3 (white) staining 3D reconstruction from 242 confocal images, z-step size=300 nm. Acquired with 60x oil NA=1.4 objective.

**Video 5 Adult rat heart gap junction staining**. Connexin 43 (red), WGA (green), and Hoechst (blue) staining 3D reconstruction from 165 confocal images, z-step size=300 nm. Acquired with 60x oil NA=1.4 objective.

**Video 6 Normal adult human kidney glomerulus.** F-actin (red) and Hoechst (green) staining 3D reconstruction from 534 confocal images, z-step size=300 nm. Acquired with 40x oil NA=1.3 objective.

**Video 7 Normal adult human kidney tiled.** F-actin (red) and Hoechst (green) staining 3D reconstruction from 253 confocal images, z-step size=300 nm. 4×4 fields of view were tiled to examine larger area. Acquired with 60x oil NA=1.4 objective.

**Video 8 Wildtype adult mouse kidney tiled.** F-actin (red) and Hoechst (green) staining 3D reconstruction from 373 confocal images, z-step size=300 nm. 3×3 fields of view were tiled to examine larger area. Acquired with 60x oil NA=1.4 objective.

**Video 9 Wildtype adult mouse liver tiled.** F-actin (white) staining 3D reconstruction from 435 confocal images, z-step size=300 nm. 3×3 fields of view were tiled to examine larger area. Acquired with 60x oil NA=1.4 objective.

**Video 10 Wildtype adult mouse liver.** F-actin (red), WGA (green), and Hoechst (blue) staining 3D reconstruction from 395 confocal images, z-step size=300 nm. Acquired with 60x oil NA=1.4 objective.

**Video 11 Wildtype adult mouse liver vasculature.** WGA (green) staining 3D reconstruction from 395 confocal images, z-step size=300 nm. Acquired with 60x oil NA=1.4 objective.

